# Environmental shedding of toxigenic *Clostridioides difficile* by asymptomatic carriers: A prospective observational study

**DOI:** 10.1101/2020.01.01.892505

**Authors:** Mayan Gilboa, Esther Houri Levi, Carmit Cohen, Ilana Tal, Carmit Rubin, Olga Feld-Simon, Adi Brom, Yehudit Eden-Friedman, Shoshi Segal, Galia Rahav, Gili Regev-Yochay, for the ShIC research group

## Abstract

**Objectives:** To compare the burden of environmental shedding of toxigenic *C. difficile* among asymptomatic carriers, *C. difficile* infected (CDI) patients and non-carriers, in an inpatient non-epidemic setting.

**Methods:** *C. difficile* carriage was determined by positive toxin-B PCR from rectal swabs of asymptomatc patients. Active CDI was defined as a positive 2-step EIA/PCR test in patients with >3 unformed stools/24 hours. *C. difficile* environmental contamination was assessed by obtaining specimens from 10 sites in the patients’ rooms. Toxigenic strains were identified by PCR. We created a contamination scale to define the overall level of room contamination that ranged from clean to heavy contamination.

**Results:** 117 rooms were screened; 70 rooms inhabited by *C. difficile* carriers, 30 rooms by active CDI patients and 17 rooms by non *C. difficile* -carriers (Control). In the carrier rooms 29 (41%) had more than residual contamination, from which 17 (24%) were heavily contaminated. In the CDI rooms 12 (40%) had more than residual contamination from which 3 (10%) were heavily contaminated, while in the control rooms, one room (6%) had more than residual contamination and none were heavily contaminated. In a multivariate analysis, the contamination score of rooms inhabited by carriers did not differ from rooms of CDI patients, yet both were significantly more contaminated than those of none carriers OR 12.23 and 11.16 (95%CI:1.5-99.96 P=0.0195, and 1.19-104.49 p=0.035), respectively.

**Conclusion:** Here we show that *C. difficile* carriers’ rooms are as contaminated as those of patients with active CDI and significantly more than those of non-carriers.

## Introduction

*Clostridioides difficile* is a leading cause of health care associated infections, resulting in significant morbidity and mortality worldwide with an estimated incidence of ~0.5 million new cases a year only in the US(1,2). This infection has severe consequences with a reported case-fatality rate of 6%–30%(3,4). The common paradigm is that transmission of this spore forming bacteria, begins when symptomatic patients with *C. difficile* infection (CDI) shed spores and contaminate their environment(5). *C. difficile* spores are highly resistant to all routine disinfectants used in hospitals(6,7). In order to reduce C. difficile transmission, infection prevention and control guidelines, including those of the CDC, ECDC, and others(8–10), recommend isolation of patients with confirmed or suspected CDI. (10,11).

The role of asymptomatic carriers in the transmission of *C. difficile* is not completely clear(12). Currently both European and American guidelines do not suggest the routine screening of asymptomatic patient or isolation of carriers (19,20). This is probably due to the fact that most studies on *C. difficile* carriers, have demonstrated relatively low levels of environmental contamination(15,17,21).

In this study, we aimed to determine the burden of environmental *C. difficile* shedding by asymptomatic carriers compared to symptomatic patients and non-carriers in an inpatient non-epidemic setting.

## Methods

### Setting

The Sheba Medical Center (SMC) is a tertiary academic medical center in central Israel with 1600-bed capacity, with 96,800 annual admissions. About a quarter of the admission are to the 300 beds of seven internal medical wards. Throughout the past two years, the overall incidence of nosocomial CDI was stable at 3.2 cases/10,000 patient-days in total and was 6.0 cases/10, 000 patient-days in the medical wards.

### Study design

This study is embedded in a larger study examining the effectiveness of *C. difficile* carriage active surveillance upon admission by rectal swabs. In this study, we assessed environmental contamination of rooms inhabited by *C. difficile* carriers, *C. difficile* infected patients and non-carriers. Between Dec 2017 and Jan 2019, a convenience sampling of rooms was performed by screening 4-5 rooms per week, where rooms occupied by the defined patients for at least 24 hours were sampled.

### Definitions of carriage and disease status

Patients admitted to an Internal Medicine ward were rectally screened for *C. difficile* carriage if they were asymptomatic and had one of the following risk factors: transferred from another institution, had a previous hospitalization during the preceding 6 months, or were older than 84 years. Carriage was determined using PCR for toxin B (tcdB) and binary Toxin(cdt) (Xpert *C. difficile*; Cepheid, Sunnyvale, CA, USA).This assay is validated and approved for testing unformed stool for CDI diagnosis,its use for screening via rectal swabbing was off-label. Non-carriers were defined as asymptomatic patients who were tested upon admission and had a negative PCR result.

Any patient with 3 unformed stools within 24 h was tested for CDI, regardless of the screening result upon admission. The 2-step EIA/PCR of unformed stool was used to define CDI(22). Glutamate dehydrogenase (GDH) antigen and toxins A and B were detected using the rapid test membrane enzyme immunoassay(EIA) (C.Diff Quik Chek Complete, Alere^MT^, Walthem, MA, USA), additionally, PCR for toxin B (Cepheid, Sunnyvale, CA, USA) was performed. CDI was defined if 2 or more of the 3 tests were positive.

### Environmental screening and environmental sample processing

In each room, 10 high-touch sites were screened, including 5 high-touch sites inside the patients’ room (floor, bedrailpatient table, armchair and call button) and 5 high-touch sites in the patients’ bathroom (floor, toilet hand-rail, toilet seat, toilet flush button and door handle). Both carriers’ room as well as CDI patients rooms are being decontaminated similarly, with hypochlorite (Actichlor plus) 5000ppm solution both on a daily basis (high touch sites only), and for terminal thorough cleaning. Rooms of non carriers are cleaned daily with 1000 PPM hypochlorite solution. As part of the protocol, the sampling was always performed in rooms inhabited for at least 24 hours by that patient and always before the daily cleaning. The samples were collected using environmental sponge-wipes (3M, St. Paul, MN, USA) applied to a designated area of each surface (5 × 20 cm). In the laboratory, the sponge-wipes were transferred to a stomacher bag (Interscience, Saint Nom, France), containing PBST (Phosphate Buffered-Saline & 0.02% Tween). The homogenized bag contents, were filtered on Brazier’s *C. difficile* selective agar (Oxoid Limited, Thermo Fisher Scientific, Perth, UK) and incubated under anaerobic conditions for 48-72 hours. Typical *C. difficile* colonies were counted and sub-cultured on Columbia-CNA agar(23). Identification was confirmed by Gram stain and VITEK®2 (Biomérieux, St.Louis, Missouri, USA) and further validated by *C. difficile* Test kit (Oxoid Limited, Thermo Scientific, Perth, UK). *C. difficile* isolation and identification was conductedat Aminolab LTD, Ness-Ziona, Israel. *C. difficile* isolates were frozen at −80°C for further analysis in our laboratory.

### Toxin B PCR assay

To verify that the environmental contamination is indeed caused by toxigenic *C. difficile*, the frozen samples were thawed and tested for Toxin B by PCR (Primers: NK104 (5’-GTGTAGCAATGAAAGTCCAAGTTTACGC-3’) and NK105 (5’-CACTTAGCTCTTTGATTGCTGCACCT-3’)) (24). Amplification was performed under the following conditions: 2 min at 95°C, followed by 35 cycles of 25s at 95°C, 35s at 54°C, 45s at 72°C and additional 5 min at 72°C.

Since the number of sites contaminated as well as the number of CFU’s both have an important role in the spread of *C. difficile*, we created a contamination scale. The scale was created a priori, following the first 10 arbitrary rooms (excluding control rooms).

The contamination scale integrated the number of contaminated sites, with the overall number of colony forming units (CFU’s) per room. Thus, designating each room a level of environmental contamination that ranged from clean to heavy contamination as shown in **Figure 1**.

**Figure 1.**
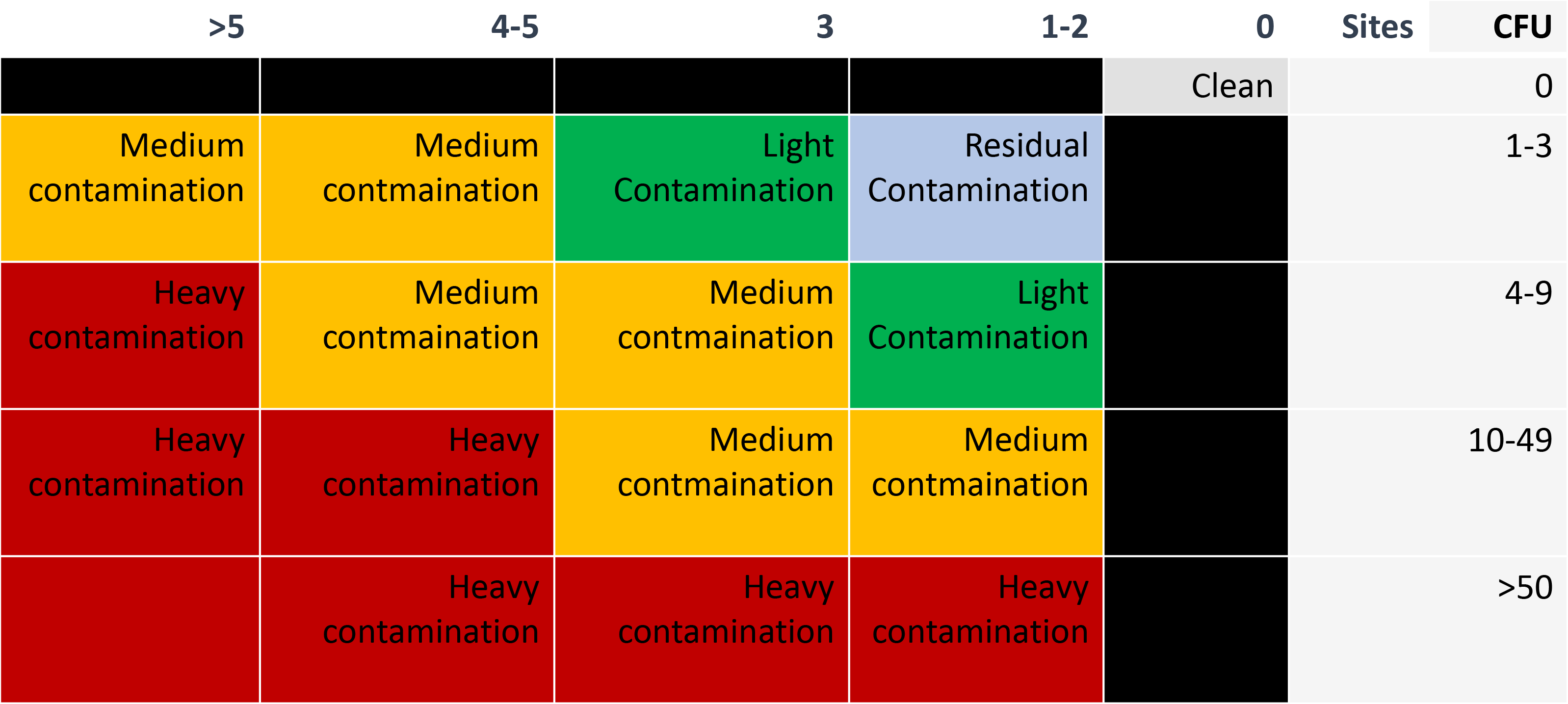
“contamination” scale- a scale integrating the number of sites contaminated with bacteria with the overall number of colony forming units (CFU’s) per site. Grey- clean, light blue- residual contamination, green- light contamination, yellow- medium contamination, red- heavy contamination

### Patient Data

Demographic and clinical data was collected from the patients’ electronic medical files, including age, gender, activity of daily life, independence status, continence status and medication use.

### Statistical analysis

Differences in patient characteristics between the 3 study groups (C. difficile carriers, non-carriers and CDI) were assessed using one way ANOVA and chi-square test when applicable. In order to compare the proportion of contaminated rooms between the 3 study groups we used chi square test. Comparison between the three groups regarding the number of colonies and sites involved was performed using non-parametric analysis of variance - Kruskal-Wallis test.

when significant differences between the three groups were indicated, post-hoc tests using Mann-Whitney tests with Bonferroni adjustment was performed.

In order to evaluate if *C. dificille* carriage is an independent predictor of room contamination after controlling for other factors, a multivariate logistic regression was applied including several characteristics of the patient inhabiting the room. Specifically, age above 84, gender, activities of daily living (ADL), antimicrobial use and *C. dificille* carriage/infection status.

A binomial variable was defined for the multivariate and univariate analysis of level of contamination (0=clean+residual; 1=>than residual).Statistical analysis were performed using SAS 9.4.

### Institutional Review Board

The study was approved by the Sheba Medical Center institutional review board. Since, only environmental swabs were taken and only few unidentified data on the patients occupying the examined rooms were collected, written informed consent was waived.

## Results

During the study period, 117 rooms were examined, in which a total of 1170 sites were screened (10 identical sites per room). Of these, 70 rooms were inhabited by asymptomatic carriers, 30 rooms by patients diagnosed with symptomatic CDI and 17 room by non-carriers. Full data could be obtained from the electronic medical records for 110 of them. Of the 1170 sites screened, in 214 (18%) *C. difficile* was isolated, the extent of contamination was diverse, from a single colony detected per site and up 200 colonies per site.

A single colony from each contaminated site was tested for the presence of toxin B by PCR and 144/214 (68%) were defined as toxigenic *C. difficile*. Only sites containing toxigenic *C. difficile* were considered contaminated.

### C. *difficile carriers’ rooms are contaminated with toxigenic* C. *difficile*

Of the 70 rooms occupied by C. *difficile* carriers, 40 (57.0%) had some degree of contamination, 29 (41.4%) had more than residual contamination, and 17 rooms (24.3%) were heavily contaminated. Average of 1.68 (±2.04) contaminated sites per room and an average of 2.93(±7.12) colonies per site. The median number of colonies was 1 (interquartile range 0-16).

In contrast, Fifteen of the 17 (88.24%) control rooms were totally clean. A single room (6%) was residually contaminated with 2 colonies detected on the bathroom floor and another room (6%) was contaminated with 11 colonies detected in 2 sites, 10 colonies on the bathroom floor and one colony on the room floor(defined as medium contaminated).

The proportion of contaminated rooms (more than residual contamination) inhabited by asymptomatic *C. difficile* carriers rooms was greater than that of rooms inhabited by non-carriers (41.4% vs. 5.9%, p=0.0057)

In the CDI group, 16 (53%) rooms had any degree of contamination, 12 (40%) rooms had more than residual contamination and 3 (10%) were heavily contaminated. An average of 1.23 (+/−2.18) contaminated sites per room, an average of 0.84 (+/−1.8) colonies per site, a median of 1 colony in each room and a interquartile range of 0-12. (**Figure 2**). Rooms inhabited by symptomatic CDI patients were more contaminated than those of non-carriers (40%vs 5.9%, p=0.02) and the proportion of a contaminated site was higher in rooms inhabited by symptomatic *C. difficile* patients than that of non-carriers (13.3% vs 1.7%, OR=20.1; 95%CI:6–67.2; p<0.0001). When comparing carrier rooms to those of symptomatic *C. difficile* patients there was no significant difference in the percentage of rooms with more than residual contamination (42% vs. 43% OR 0.8 p=0.63 CI 0.64-1.91)

**Figure 2.**
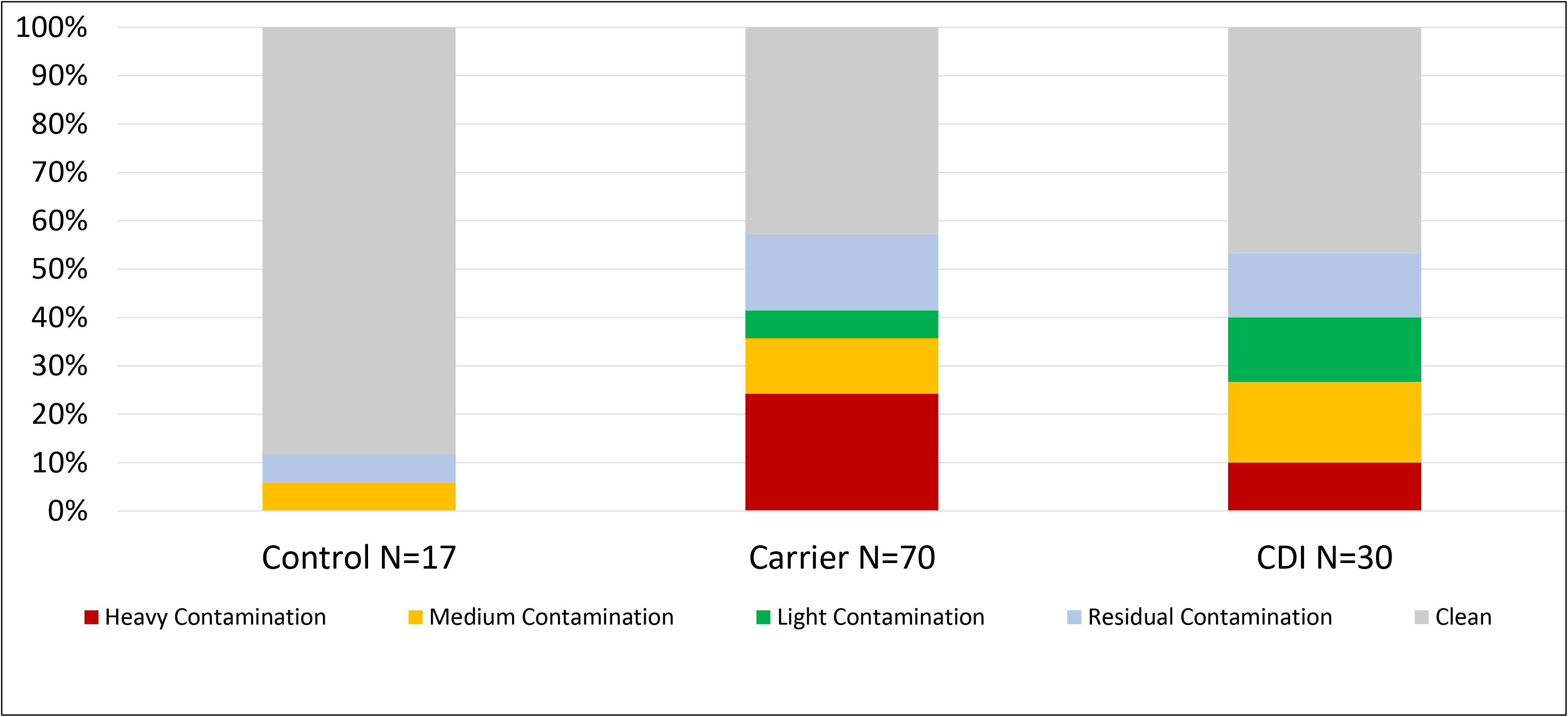
Environmental contamination of rooms occupied by CDI, carrier and non-carrier patients. Grey- clean, light blue- residual contamination, green- light contamination, yellow- medium contamination, red- heavy contamination

### Contamination Distribution among the various sites tested

The most contaminated sites in patient’s environment were the floors with 19 (27%) positive samples of floors from carrier rooms and 7 (23%) in the CDI group. Even the very rare contamination we observed in non-carriers was of the floor. The bathroom floors were also highly contaminated, with 15 (21%) of the carriers’ rooms and in 5 (16%) of the CDI patients’ rooms. Additionally, 19 (27.1%) and 5 (16.67%) bedrails where contaminated in carrier and CDI rooms, respectively. Armchairs were contaminated in 13 (19%) and 4 (13.33%) of carrier and CDI rooms, respectively. There was no significant statistical difference in the proportion of contamination for each site between carriers and CDI patients (**Figure 3**).

**Figure 3.**
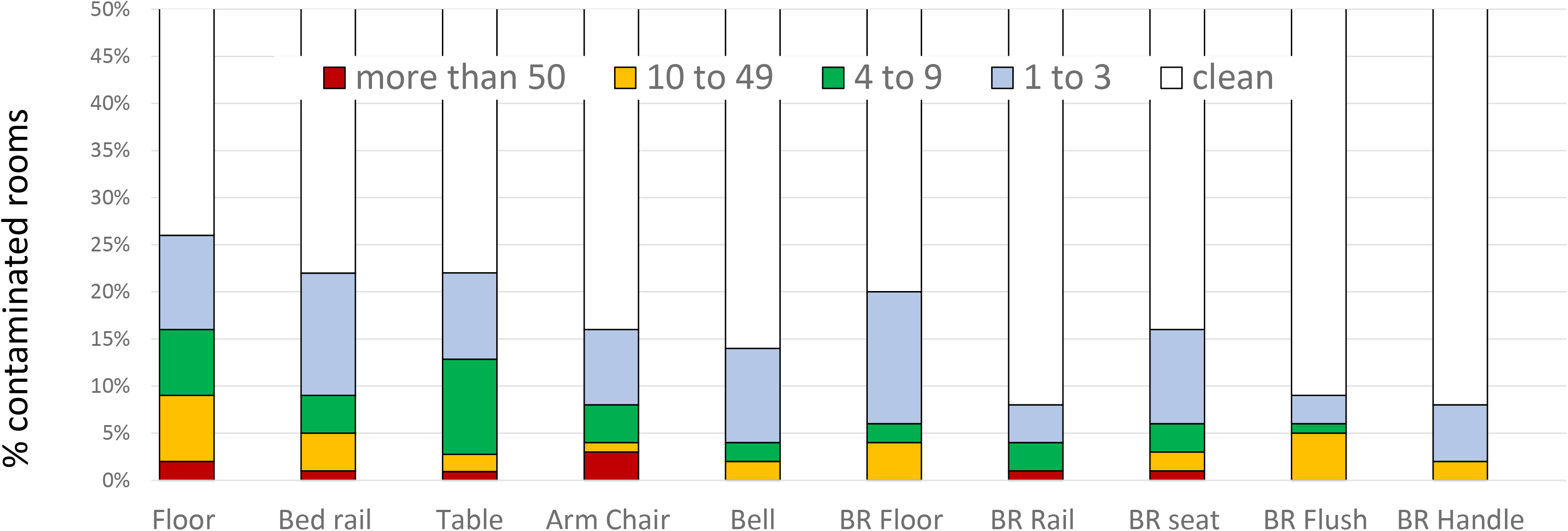
Environmental contamination of 10 high-touch points. Grey represents no CFU’s, light blue- 1-3 CFU’s, green-4-9 yellow-10-49 red-above 50

### Patient Characteristics

To determine whether carriage status of the room inhabitant is a true independent predictor of contamination, we initially compared patient characteristics, of the 3 study groups: asymptomatic C. *difficile* carriers, symptomatic CDI patients and non- *C. difficile* carrier controls. Indeed, we found that the groups differed in a few parameters: Age, the CDI group was significantly younger than that of the carrier group (mean age 60.7 (±17.3) as compared to 70.3 (±15.3), only 7% of CDI patients were older than 84 as compared to 14% of the carriers (p=0.02). Non-CDI directed antimicrobial agent use was significantly lower in the CDI group, where only 24% received antimicrobial agents as compared to 49% of the carriers and 71% of non-carriers (p<0.01) (**Table 1**). The proportion of patients treated with anti-*C. difficile* directed antibiotics on the day of environmental screening was significantly higher among CDI patients vs. carriers and non-carriers (80.8%, 7.5% and 0%, respectively). Counter to our guidelines, five *C. difficile* carrier patients were treated with metronidazole as a preventive measure while on wide spectrum antibiotic treatment (for another infectious disease).

**Table 1.** patient population

|  |  | CD-Non Carriers<br>N=17 |  |  |  | CD Carriers<br>N=70 |  |  |  | Active CDI<br>N=30 |  |  |  | P<br>value |
| --- | --- | --- | --- | --- | --- | --- | --- | --- | --- | --- | --- | --- | --- | --- |
|  |  | % | N | Mean | SD | % | N | Mean | SD | % | N | Mean | SD |  |
| **age** |  |  |  | 74.1 | 14.3 |  |  | 70.3 | 15.3 |  |  | 60.7 | 17.3 |  |
| **Gender** | *male* | 64.7% | 11 |  |  | 52.2% | 36 |  |  | 64.3% | 18 |  |  | 0.43 |
| **Incontinence** | *Incontinent* | 70.6% | 12 |  |  | 56.7% | 38 |  |  | 46.2% | 12 |  |  | 0.29 |
| **PPI use** | *PPI* | 35.3% | 6 |  |  | 35.8% | 24 |  |  | 42.3% | 11 |  |  | 0.91 |
| **Any**<br>**antibiotic usage\*** | *antibiotics* | 70.6% | 12 |  |  | 49.3% | 33 |  |  | 20% | 6 |  |  | 0.029 |
| **Anti CD**<br>**treatment\*\*** |  | 0.0% | 0 |  |  | 7.5% | 5 |  |  | 80.8% | 21 |  |  | <0.001 |

\*Any antibiotic usage- excluding metronidazole IV or PO and Vancomycin PO
\*\*Anti CD treatment- anti *C. difficile* antibiotics (Metronidazole PO or IV and Vancomycin PO)).

**Table 2:** Multivariate analysis of predictors for environmental contamination

|  |  |  | 95% CI |  |  |
| --- | --- | --- | --- | --- | --- |
| variable |  | OR |  |  | p value |
| **Age** | **older then 84 vs**<br>**younger then 84** | 1.07 | 0.334 | 3.478 | 0.9008 |
| **Gender** | **Female Vs Male** | 0.744 | 0.319 | 1.733 | 0.4931 |
| **Dependency In ADL**<br>**activites** | **Dependent Vs**<br>**Indpendent** | 0.595 | 0.255 | 1.389 | 0.2297 |
| **Antibiotics usage** | **yes vs no** | 1.005 | 0.421 | 2.401 | 0.9913 |
| **PPI** | **yes vs no** | 0.87 | 0.372 | 2.034 | 0.7481 |
| **Carriage Status** | **carrier vs control** | 12.5 | 1.5 | 102.7 | 0.0186 |

**Table 3.**
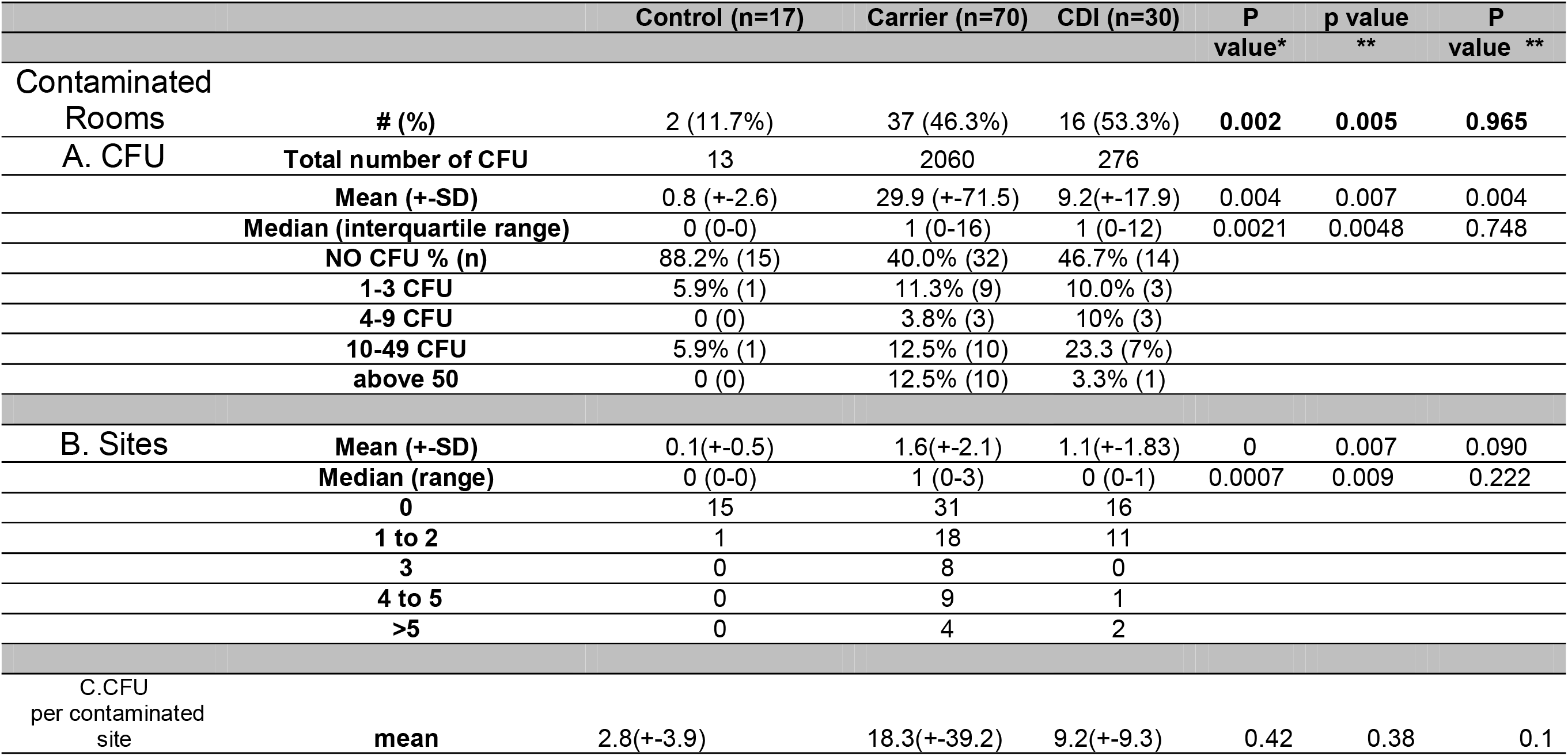
A. Total number of colony forming units (CFU) in all rooms, Mean number of CFU per participant’s room, Median number of CFU’s per participant room and distribution of amount of CFU per room. B. Mean number of contaminated sites per participants room, Median number of contaminated sites per participants room and distribution of number of contaminated areas. C. Mean number of CFU per contaminated site. * p value between carrier and control ** p value between CDI and control *** p value between carrier and CDI

To adjust for these differences and determine whether carriers’ rooms are independently associated with higher environmental contamination, we conducted a multivariate analysis correcting for age (older and younger of 84), gender, dependency in ADL activities, PPI usage and non CDI specific antibiotics usage. After adjustment, a carrier room was a significant independent predictor for contamination (more than residual contamination)with OR=10.8 (95%CI:1.33-87.95, p=0.026). Similarly, the odd ratio of a CDI patient room was 11.16 (95%CI: span style=“font-family:Calibri”>1.19-104.49, p=0.0345.

## Discussion

The role of *C. difficile* carriers in *C. difficile* transmission is controversial. Current guidelines of different societies and organizations do not recommend screening patients for *C. difficile* carriage, and *C. difficile* carriers remain unrecognized and are not isolated. . To date, data suggesting that asymptomatic *C. difficile* carriers are a source of spread of *C. difficile* in the environment have only been in frequently published ^(13,15–18)^.

Here we showed that the environment of asymptomatic *C. difficile* carriers is as contaminated as that of symptomatic CDI patients. We found toxigenic bacteria throughout the carriers’ environment in various sites of the patients rooms and attached bathrooms.

Previous studies based on genetic sequencing such as multilocus sequence typing (MLST) and multilocus variable-number tandem-repeat analysis (MLVA) found that only 25%− 55% of symptomatic infections could be linked to a previously identified CDI patient(13,27). Asymptomatic carriers were suggested as one of the possible sources and reservoirs

Longtin et al. have reported an intervention study, where C.*difficile* carriers were detected and partially isolated(18) and showed that this resulted in decrease in CDI incidence. Yet, it was unclear whether this was due to better antibiotic stewardship of *C. difficile* carriers or due to less transmission by carriers.

Of the 10 different high-touch sites assessed, the most contaminated site was the floor, this was true for both the bedroom and the bathroom. Hospital floors are frequently contaminated with pathogens and it has been previously established that floors are important reservoirs of bacteria in the patient’s environment. (29). This finding emphasizes the importance of particular focus on floor cleaning in hospitals. Several previous studies have shown that asymptomatic C. *difficile* carriers can contaminate the environment but these studies differed from ours in several aspects (16,20,29–30). Our study is the first study in a non-epidemic hospital setting that screened a substantial number of asymptomatic patients for C. *difficile* carriage and examined numerous sites in their rooms.

Our study has several limitations. We could not determine the genetic identity of the environmental strain to that of the patient occupying that room. Thus, one could argue that the contamination couldave been from previous CDI patients occupying that room. Yet, the fact that rooms of non-carriers where practically clean, strongly suggests that the environmental shedding was not incidental and was related to the carriage status of the patient. We also tested only a single colony by PCR, to define environmental contamination by toxigenic *C. difficile*. Therefore, in a case where there were multiple concomitant clones this could potentially cause underdetection of toxigenic strains. Interestingly in the control group 100% of isolated colonies were toxigenic, if there was such an underestimation it could have been only in the carrier or CDI group where 77% and 55% of isolated colonies where toxigenic respectively.

The carrier group in our study differed significantly from the CDI group, in both age and antibiotics usage. An explanation for this could be that one of the criteria for screening asymptomatic patients upon admission was age above 84. The fact that antibiotic coverage differed significantly between the three groups reflects the practice of withholding antibiotic treatment in patients with active CDI infection, and also that information about carriage status could have encouraged medical staff to use antibiotics more cautiously in this population. To overcome this limitation we conducted a multivariate analysis that showed that even after adjusting for these variables the rooms or carriers, as well as of CDI patients were significantly (~10 times) more contaminated than non-carriers.

Last, rooms were screened at a single time-point, which differed between patients, but was >24 hours of hospitalization, when most of the CDI patients were already treated with anti- *C. difficile* antibiotics. This could cause under-detection of overall *C. difficile* contamination in rooms occupied by CDI patients (16), which may actually be more contaminated than carriers’ rooms, as other studies have reported (15,26).

In conclusion our study suggests that a major source for *C. difficile* transmission in the hospital, is probably by unidentified *C. difficile* carriers, which are not routinely isolated, they spread *C. difficile* to their environment, which is not routinely disinfected with sodium hypochlorite. Our study adds to accumulating data supporting the need to screen asymptomatic patients, detect *C. difficile* carriers and address them similarly as to CDI patients, both in isolation and cleaning practices. It is yet to be shown that obtaining these measures will in fact reduce the rates of *C. difficile* infections in hospitals. Further studies are required to demonstrate the efficacy of detecting *C. difficile* carriers and limiting their environmental contamination on reducing CDI incidence.

## Acknowledgments

The SHIC research group: Howard Amital, Sharon Beni, Ilan Ben-Zvi, Natasha Blausov, Adi Brom, Carmit Cohen, Olga Feld-Simon, Ronen Fluss, Mayan Gilboa, Yehudit Eden-Friedman, Shiraz Halevy, Esther Houri-Levi, Amit Hupert, Amnah Jbarien, Naty Keller, Avshalom Leibowitz, Haim Mayan, Leonid Maizels, Eyal Meltzer, Nani Pinas-Zade, Galia Rahav, Shir Raibman-Spector, Gili Regev-Yochay (PI), Marina Romiantsev, Dalit Shachar, Kassem Sharif, Gadi Segal, Shoshi Segal, Amitai Segev, Gill Smollan, Shmuel Stienlauf, Ilana Tal, Hagit Yonath, Tal Zilberman-Daniels, Eyal Zimlichman. We greatly acknowledge the work and assistance of the whole Infection Prevention & Control team at the Sheba medical Center, Ms. Efrat Steinberger for coordinating the study and The Sheba medical Center management for their full support. Funding: This study was funded by Magen–Chaim (Life-Shield) grant.

Disclosure- Dr. Gilboa, Dr. Houri Levi, Dr. Cohen, Mrs. Tal, Dr. Rubin, Dr. Feld-Simon, Dr. Brom, Dr. Eden- Friedman, Mrs. Segal, Prof. Rahav, and Prof. Regev-Yochay have nothing to disclose.

